# A novel patient-derived cellular platform for validating microglia-targeted therapeutics for Alzheimer’s disease

**DOI:** 10.1101/2023.08.17.552618

**Authors:** Carla Cuní-López, Romal Stewart, Satomi Okano, Garry L. Redlich, Mark W. Appleby, Anthony R. White, Hazel Quek

**Author notes:** Contributed equally (Anthony R. White, Hazel Quek). Correspondence: Carla Cuní-López, Anthony R. White, Hazel Quek. **Email:**.

## Abstract

The scarcity of effective biomarkers and therapeutic strategies for predicting disease onset and progression in Alzheimer’s disease (AD) is a major challenge to improve much-needed therapeutic outcomes. Conventional drug discovery approaches have been unsuccessful in providing efficient interventions due to their ‘one-size-fits-all’ nature. As an alternative, personalised drug development holds promise to pre-select responders and identify suitable drug efficacy indicators. In this study, we established a preclinical drug testing strategy by assessing the efficacy of anti-inflammatory drugs in 2D and 3D *in vitro* models of monocyte-derived microglia-like cells (MDMi) derived from AD and mild cognitive impairment (MCI) patients, and matched healthy individuals. We observed that the cytokine inflammatory profiles of MDMi in response to drugs clustered separately between cohorts, with the 3D model showing a more defined separation between healthy and patient donors than 2D. By ranking donor and cytokine responses to drugs, we identified that drug efficacy was limited in AD patients and involved cohort-specific responsive cytokines. Our findings suggest that MDMi models have the potential to predict disease progression, stratify responders and identify biomarkers for estimating the efficacy of microglia-targeted drugs. Together, our pipeline could serve as a valuable tool to enhance the clinical translational value of preclinical drug screens and ultimately improve drug outcomes for AD.

## Introduction

With Alzheimer’s disease (AD) and related dementias becoming increasingly prevalent, there is a pressing medical need to develop effective therapeutic strategies. Approved interventions in recent years, such as the controversial anti-amyloid drug aducanumab, have been extremely scarce and their efficacy in providing cognitive benefits is being questioned (1). These minimal drug approval rates can be attributed to the complex molecular disease drivers, inaccurate diagnostic criteria, and inefficient biomarkers of disease onset and progression (2, 3).

Current clinical testing approaches for assessing new therapeutics in AD also come with important caveats. These challenges include the variability in treatment responses to drug and placebo among participants, the use of inappropriate statistical methodologies to assess drug effect size, and a lack of correlation between biomarker changes and cognitive improvement (1, 4). Furthermore, ineffective preclinical AD drug efficacy studies contribute to a flawed drug development pipeline given their poor rigour, reproducibility, predictive value and limited translation into clinical outcomes. These limitations are in part attributed to the shortcomings of currently used preclinical model systems, including animal and cell disease models, which inadequately recapitulate human disease processes (5). To address these challenges, there is a need for more informative preclinical studies that differentiate between asymptomatic individuals and patients at various disease stages. Such studies may lead to an improved subject selection in subsequent clinical trials and the identification of more predictive drug efficacy targets with translational relevance to patients.

Novel technologies have emerged to improve preclinical drug testing for AD. One of these tools is the generation of the monocyte-derived microglia-like cell model (MDMi) (6). MDMi offer several advantages over other human microglia cell models, such as their cost-effective and rapid derivation from easily accessible blood samples, and have shown improved characteristics when differentiated in a more physiologically relevant 3D model (7, 8). Furthermore, MDMi permit the longitudinal assessment of patient-specific drug responses, allowing for the simultaneous monitoring of drug effects in patients (9). Additionally, MDMi exhibit disease-specific features associated with neurodegenerative diseases, which validates their use as an *in vitro* platform for preclinical drug studies targeting neuroinflammation (10–13).

Here, we present a novel drug testing pipeline that uses 2D and 3D MDMi models derived from healthy donors, and patients with AD and mild cognitive impairment (MCI). Our approach uses multidimensional analysis to select responders and identify the most informative targets of drug efficacy. We validated this pipeline with two FDA-approved drugs: minocycline and IC14. Minocycline, a well-known anti-inflammatory drug, has shown effectiveness in rodent and immortalised microglial models but has failed in clinical testing for AD patients (14–16). IC14, an anti-CD14 monoclonal antibody, is currently under clinical investigation for treating inflammation related to SARS-Cov-2 and amyotrophic lateral sclerosis (ALS) and has potential for repurposing in AD patients (17–19). We have identified differences in MDMi responses to both drugs, which could aid in predicting the efficacy of new compounds before clinical evaluation. Overall, this pipeline holds promise for pre-selecting patients along with validating and predicting candidate drugs before clinical trials, and could yield informative biomarkers of drug efficacy in a personalised manner.

## Results

### MDMi characterisation and applicability for drug testing

MDMi in both 2D and 3D cultures exhibited characteristics of microglia-like cells, including a signature ramified morphology and protein expression of IBA1 (**Fig. 1A**). Moreover, our previous study demonstrated that in 3D, both the microglia-like signature and the expression of disease-related alterations in AD patient-derived MDMi were enhanced compared to 2D cultures (12). These findings suggest that MDMi can serve as a personalised cell model for studying drug response profiles in a patient-specific manner.

**Fig. 1.**
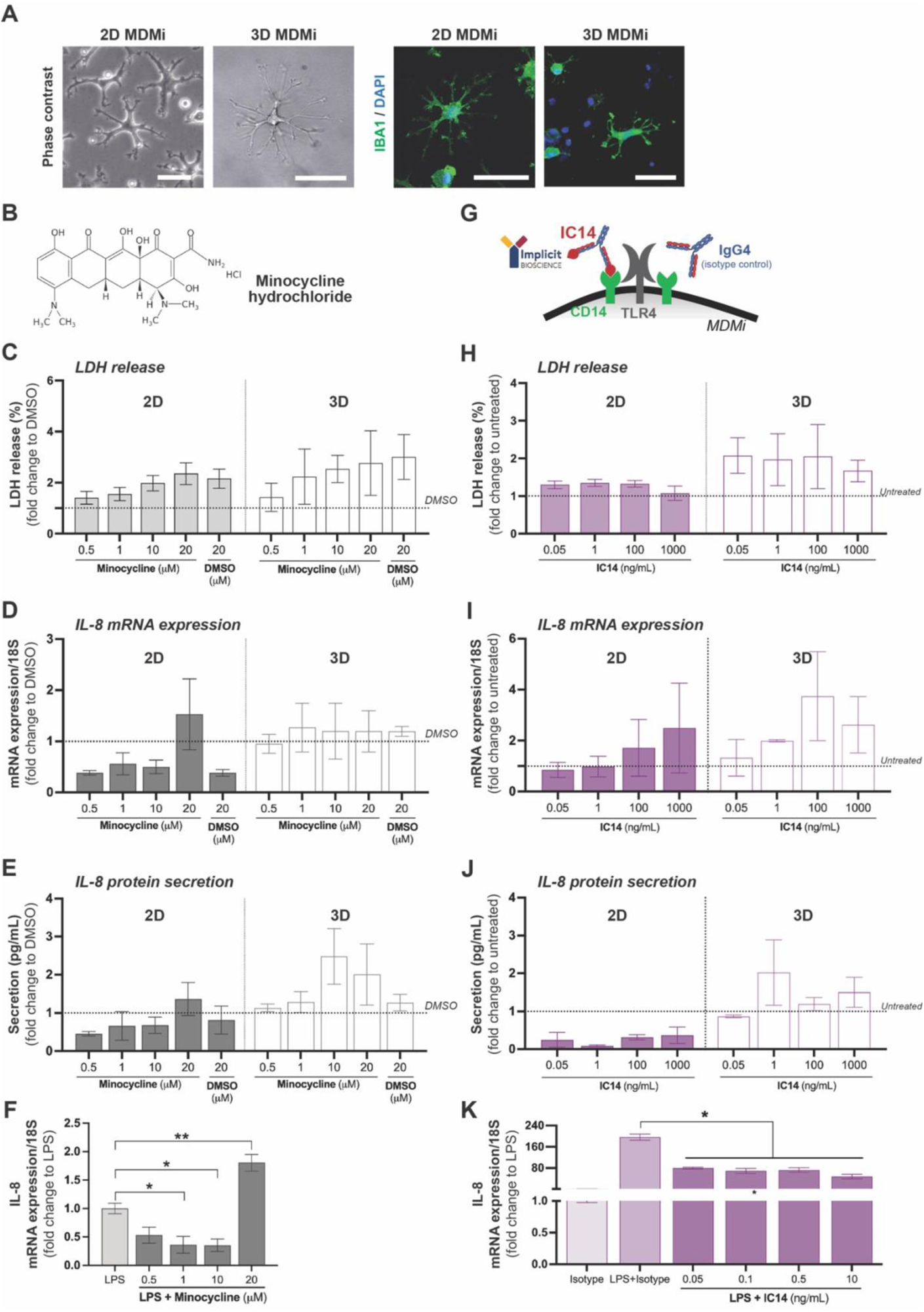
MDMi characterisation and assessment of drug cytotoxicity and anti-inflammatory capacity. **(A)** Phase contrast and immunofluorescence images of 2D and 3D MDMi derived from young healthy donors. Scale bars are 50 μm. **(B)** Chemical structure of minocycline hydrochloride. Minocycline-mediated toxicity in MDMi measured by **(C)** LDH release, **(D)** *IL-8* mRNA expression and **(E)** IL-8 protein secretion (all calculated as fold change to DMSO 1 μM, represented with dotted lines). **(F)** Anti-inflammatory effects of minocycline against LPS measured by *IL-8* mRNA expression (calculated as fold change to LPS alone treatment). **(G)** Schematic representation of IC14 and isotype control (IgG4) antibodies targeting the CD14 receptor on the membrane surface of MDMi. IC14-medaited toxicity in MDMi was measured by **(H)** LDH release, **(I)** *IL-8* mRNA expression and **(J)** IL-8 protein secretion (all calculated as fold change to untreated, represented with dotted lines). **(K)** Anti-inflammatory effects of IC14 against LPS measured by *IL-8* mRNA expression (calculated as fold change to LPS alone treatment). One-way ANOVA in **C-F**, **H-K**; \**P* < 0.05. Data are presented as mean ± SEM and include 3 donors (*n* = 3).

### Optimisation of minocycline and IC14 doses in MDMi from young healthy donors

Minocycline and IC14 working doses were first optimised in MDMi from young healthy control (HC) donors (<50 years old) (**Table 1**).

**Table 1.** Summary of young healthy donor information for drug optimisation studies.

|  |  | Young Healthy donors |
| --- | --- | --- |
| N° |  | *n* = 3 |
| Sex | Females (%) | 0% (0/3) |
|  | Males (%) | 100% (3/3) |
| Age (mean ± SD) |  | 28.3 ± 6.8 |

SD = standard deviation

For minocycline hydrochloride (**Fig. 1B**), MDMi were treated with a range of concentrations (0.5, 1, 10 and 20 μM) or vehicle (DMSO) for 24h, and cell viability was then measured by lactate dehydrogenase (LDH) release and IL-8 at both mRNA expression and protein secretion levels. We observed a dose-dependent increase in LDH release in both 2D and 3D models (**Fig. 1C**). Furthermore, *IL-8* mRNA expression levels were upregulated at 20 μM in 2D with minimal changes observed in 3D (**Fig. 1D**). Protein secretion of IL-8 increased at 20 μM in 2D and at 10 μM and 20 μM in 3D (**Fig. 1E**), indicating that high doses of minocycline (10 - 20 μM) induce a toxic pro-inflammatory response in MDMi. To examine the anti-inflammatory effect of minocycline, 2D MDMi were stimulated with lipopolysaccharide (LPS) followed by treatment with increasing minocycline concentrations (0.5, 1, 10, and 20 μM). We observed a significantly reduced *IL-8* mRNA expression in 2D MDMi at 1 μM and 10 μM (**Fig. 1F**). Therefore, we selected a dose of 1 μM for subsequent studies due to its non-toxicity and anti-inflammatory effects.

IC14, a recombinant chimeric monoclonal antibody targeting CD14, is a key receptor found in microglia and other immune cells involved in mediating inflammatory responses (20). IgG4 was used as isotype control to assess the specificity of IC14-mediated effects in MDMi (**Fig. 1G**). To determine an optimal dose of IC14, young HC MDMi were treated with a range of concentrations (0.05, 1, 100 and 1000 ng/ml) for 24h and subsequently cell viability and anti-inflammatory efficacy were assessed by examining LDH release and IL-8 response, respectively. LDH showed similar release levels across all concentrations in both platforms, indicating no significant cytotoxicity (**Fig. 1H**). However, *IL-8* mRNA expression exhibited a dose-dependent upregulation in both 2D and 3D models (**Fig. 1I**) with no changes observed at the protein secretion level (**Fig. 1J**), thus suggesting increased toxicity at higher doses. The efficacy of IC14 against LPS-induced inflammation was then evaluated in 2D MDMi by quantifying *IL-8* mRNA expression. Our results showed downregulated *IL-8* levels at all tested IC14 concentrations compared to isotype control (**Fig. 1J**). Based on these findings, a dose of 0.05 ng/mL was selected for further experiments due to its minimal toxicity and anti-inflammatory efficacy.

### Unmasking responsive cytokines and responders: overcoming limitations of cohort-based drug efficacy analysis

To investigate the responses to minocycline and IC14 in MDMi derived from dementia patients and matched healthy individuals, we established 2D and 3D MDMi models using a cohort of HC (*n*=6), mild cognitive impairment (MCI) (*n*=3), and AD patients (*n*=3), as outlined in **Fig. 2A**. HC individuals were matched for age, sex and apolipoprotein E (*APOE*) genotype with the patient cohorts and were recruited based on their genetic risk to develop late-onset AD (21). Donor demographic and clinical details are provided in **Table 2**. Inflammatory profiles were characterised following LPS exposure and drug treatments by analysing cytokine responses at both mRNA expression and protein secretion levels.

**Fig. 2.**
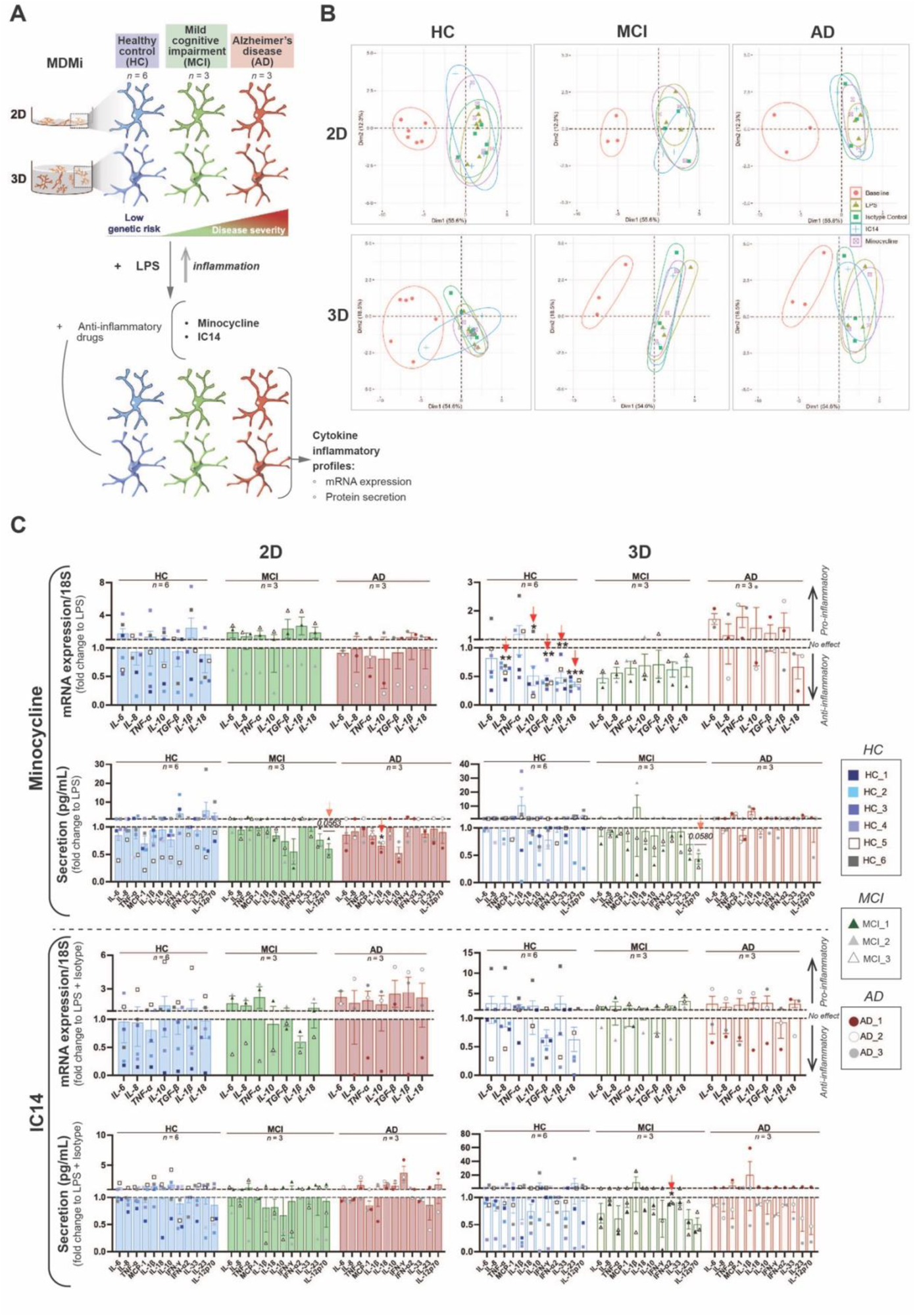
Overall scheme of the study and drug efficacy analysis in the study cohorts. **(A)** MDMi from three cohorts of donors (healthy control, HC; mild cognitive impairment, MCI; and Alzheimer’s disease, AD) were generated in 2D and 3D models. Following induction of inflammation with LPS, the cytokine inflammatory profiles of MDMi in response to drugs (*i.e.,* minocycline 1 μM and IC14 0.05 ng/mL) were characterised at the mRNA expression and protein secretion levels. These profiles were then compared between donors using multidimensional analyses. **(B)** Principal component analysis (PCA) plots showing the clustering of cohort-specific MDMi inflammatory profiles in untreated (baseline), after LPS alone (LPS) or after LPS with drugs (minocycline, IC14, isotype control) treatments. **(C)** Cytokine responses (mRNA includes *n* = 7 cytokines; and secretion includes *n* = 12 cytokines) to minocycline and IC14 treatments separated by cohort. Values were calculated as fold change to LPS (for minocycline) or LPS with isotype (for IC14), represented with dotted lines. Responses above and below the fold change indicate pro-inflammatory or anti-inflammatory effect, respectively. One-sample *t-*test performed on log-transformed data in **C**; \**P* < 0.05; \*\**P* < 0.01, \*\*\**P* < 0.001 (red arrows). Data are presented as mean ± SEM and include 6 donors (*n* = 6) in HC, 3 donors (*n* = 3) in MCI; and 3 donors (*n* = 3) in AD.

**Table 2.** Demographic and clinical details of patient and healthy control donor cohorts.

|  | **ID** | **Age**<br>**(years**<br>**old)** | **Sex** | **APOE**<br>**genotype** | **Amyloid-**<br>**β burden** | **Centiloid**<br>**amyloid-β**<br>**PET scale** | **Clinical**<br>**presentation** |
| --- | --- | --- | --- | --- | --- | --- | --- |
| **Healthy**<br>**Control** | HC_1 | 55 | Male | ε3/3 | Low | -7.53 | Asymptomatic |
|  | HC_2 | 50 | Female | ε3/3 | Low | -3.75 | Asymptomatic |
|  | HC_3 | 67 | Female | ε3/3 | Low | -4.89 | Asymptomatic |
|  | HC_4 | 64 | Male | ε3/3 | Low | -2.85 | Asymptomatic |
|  | HC_5 | 63 | Female | ε3/4 | Low | -4.53 | Asymptomatic |
|  | HC_6 | 58 | Male | ε3/3 | Low | -6.42 | Asymptomatic |
| **Mild**<br>**cognitive**<br>**impairment** | MCI_1 | 68 | Male | ε3/4 | High | unknown | Amnestic<br>multiple-<br>domain |
|  | MCI_2 | 76 | Female | ε3/3 | Very high | unknown | Amnestic<br>multiple-<br>domain |
|  | MCI_3 | 70 | Male | ε3/3 | Low | -9.4 | Amnestic<br>multiple-<br>domain |
| **Alzheimer's**<br>**disease** | AD_1 | 70 | Female | ε4/4 | Very high | 93.48 | Amnestic<br>single-domain |
|  | AD_2 | 66 | Male | ε3/3 | Very high | 116.75 | Amnestic |
|  | AD_3 | 55 | Female | ε3/3 | Very high | 96.16 | Amnestic |

APOE: apolipoprotein E; PET: positron emission tomography

A principal component analysis (PCA) of MDMi inflammatory profiles revealed a clear distinction between baseline (untreated) and post-LPS or drug treatment conditions across all cohorts and models (**Fig. 2B**). Baseline responses in HC showed higher overlap with other treatments, likely due to high inter individual variability (**Fig. 2B**). These findings confirm that MDMi from both healthy and patient donors respond to LPS stimulation and subsequent minocycline and IC14 or isotype control treatments.

We next evaluated the efficacy of minocycline and IC14 in reducing LPS-induced inflammation by using a conventional drug efficacy analysis that involves averaging cohort responses (**Fig. 2C**). This analysis revealed limited anti-inflammatory drug efficacy, with only certain responses and cohorts showing significant effects (red arrows in **Fig. 2C**). These results suggest that the observed minimal efficacy of minocycline and IC14 in reducing LPS-induced inflammation could be attributed to substantial intra-cohort heterogeneity, which may mask the presence of selective responders (*e.g.*, MCI_3 in 2D mRNA for minocycline; AD_3 in 3D secretion for IC14) (**Fig. 2C**). Moreover, inconsistencies in drug responses between 2D and 3D MDMi models suggest that the *in vitro* extracellular environment strongly influences MDMi physiology.

Our results demonstrate that a cohort-based analysis approach offers a low-resolution representation for evaluating drug efficacy and comparing individuals with different disease status, thus limiting the utility of MDMi models as platforms to discriminate between patients with varying disease severities and stratify responders.

### Donor-specific multidimensional analysis reveals heterogeneity of drug responses and discrimination between healthy and patient inflammatory profiles

To enhance the translational potential of preclinical drug outcomes in dementia patients using MDMi models, we assessed individual cytokine inflammatory profiles across all donors in the cohorts. Heatmaps of these profiles display specific cytokines measured as mRNA expression or protein secretion readouts in 2D or 3D model systems (**Fig. 3**, **Fig. S1**). We found that baseline cytokine levels were consistently lower in both 2D and 3D models for all donors as compared to LPS- and drug-treated MDMi (**Fig. S1A-B**). Notably, hierarchical clustering and PCA of baseline cytokine profiles revealed a more distinctive separation between healthy and patient cohorts in 3D models than in 2D (**Fig. S1C**), suggesting that 3D MDMi responses may have greater predictive power for selecting individuals prior to drug evaluation.

**Fig. 3.**
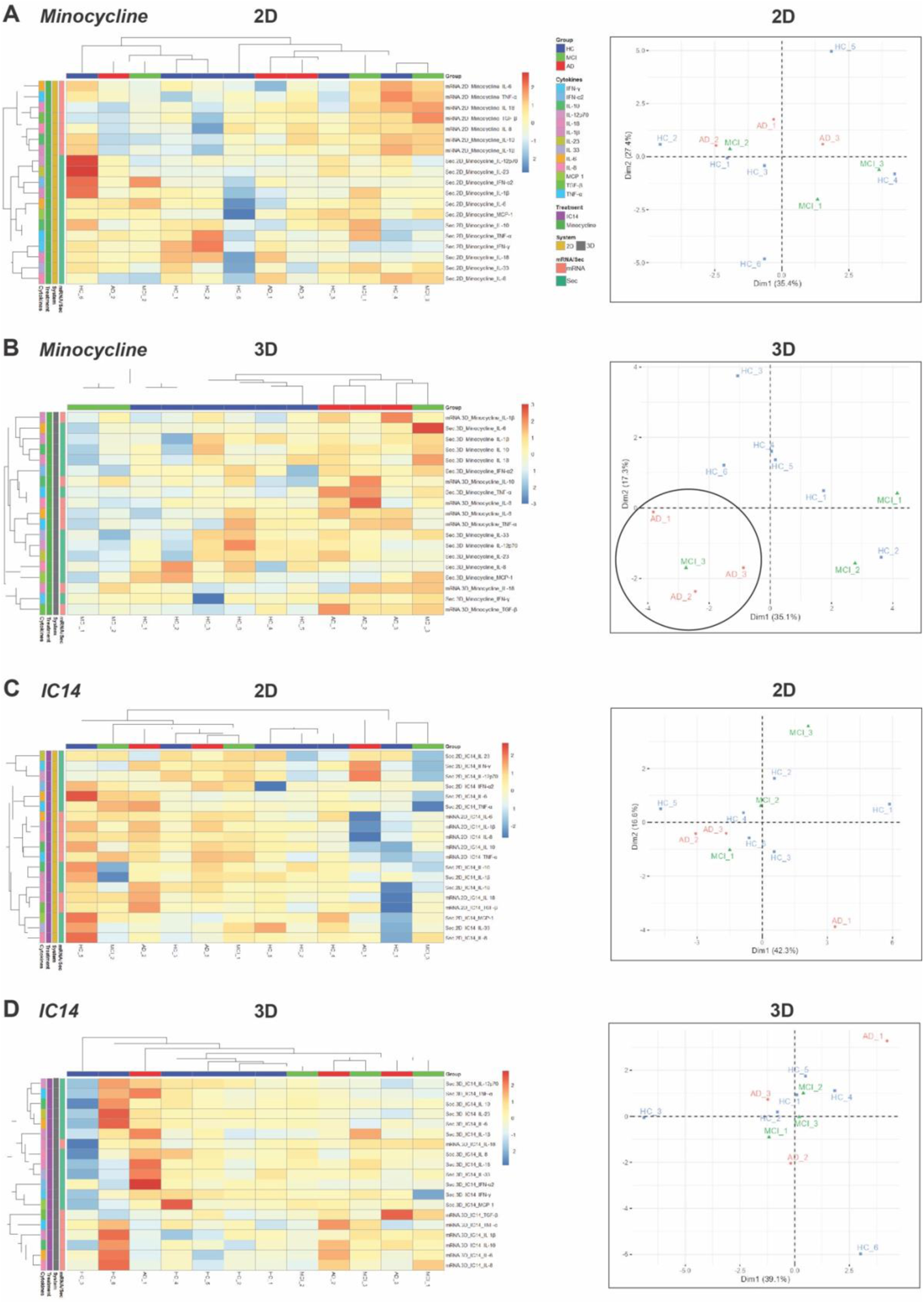
Heatmap representation of donor-specific drug response inflammatory profiles and comparison using hierarchical clustering and PCA. Heatmaps of cytokine responses (mRNA and secretion) in HC, MCI and AD donors with hierarchical clustering of donor profiles (*n* = 12) and cytokine responses (total *n* = 19, which includes *n* = 12 cytokines for secretion analyses and *n* = 7 cytokines for mRNA analyses) following treatment with minocycline in **(A)** 2D and **(B)** 3D models; and IC14 in **(C)** 2D and **(D)** 3D models. Heatmap values were normalised to LPS (minocycline) or LPS with isotype (IC14), log-transformed and scaled. PCA analyses, presented as plots on the right-hand side, included all normalised cytokine response values (*n* = 19) from each donor.

Comparison of cytokine profiles in response to minocycline and IC14 treatments demonstrated variability among donors within the same cohort. In 2D models, drug responses exhibited stronger similarities between donors from different cohorts (particularly HC and patients) than between donors within the same cohort (**Fig. 3A, C**). In contrast, 3D models showed distinct clustering of patient profiles separate from healthy donor profiles (**Fig. 3B, D**). Notably, the clustering of 3D patient responses in PCA plots was less evident for IC14 compared to minocycline (**Fig. 3B, D**) due to increased inter-patient variability in IC14 responses. A Kruskal-Wallis test identified *TGF-β* and *IL-18* mRNA expression in 3D models for minocycline and IC14, respectively, as the most significant (*P*<0.05) drug responses differentiating cohorts (**Table S1, Fig. S2**). Other notable differential responses (0.05<*P*<0.1) included IL-12p70 secretion (2D, minocycline), *IFN-γ* and *IL-18* (2D, IC14), and *IL-6* mRNA expression (3D, minocycline) (**Fig. S2**).

### Approach for identifying inflammatory biomarkers with predictive drug efficacy potential

To elucidate the most responsive inflammatory targets in the study cohorts, we ranked the means of the analysed responses based on cytokine type, readout, and model system (**Fig. 4**). Top-ranked responses were selected based on a minimum 20% reduction compared to LPS or LPS and isotype control treatments and indicated anti-inflammatory drug efficacy. The remaining responses displayed either a lack of modulation or increased inflammation (pro-inflammatory action) (**Fig. 4A, D**; **Supplementary data 1, 2**), suggesting a potential lack of drug efficacy or adverse effects. The selection of a 20% cut-off was based on the results of a randomised controlled trial that showed an average 20% reduction of IFN-γ and IL-10 plasma levels in depression patients following treatment with minocycline for 4 weeks (22).

**Fig. 4.**
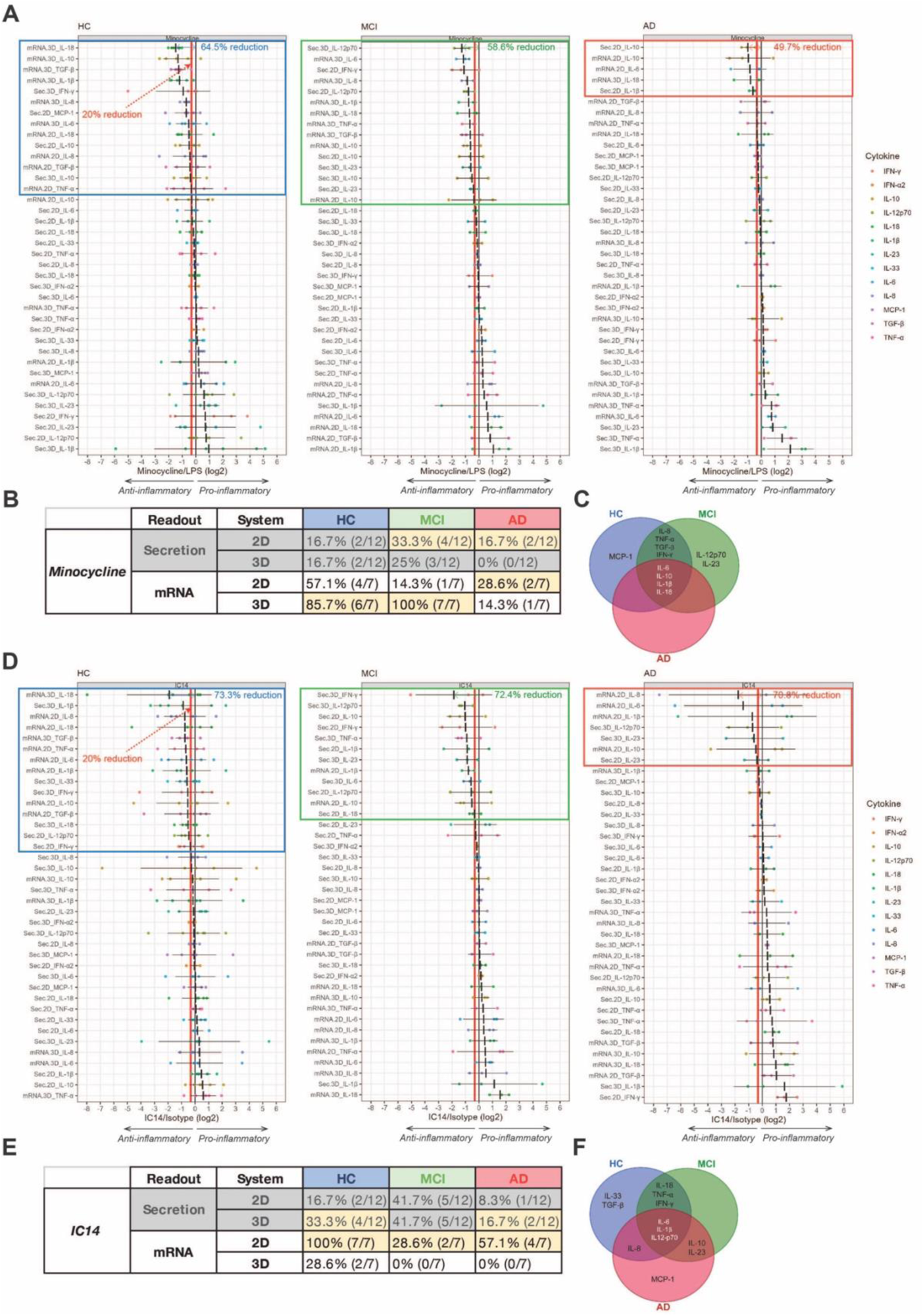
Ranking of cytokine responses to drugs. **(A, D)** Minocycline and IC14 responses (total of *n* = 38, which includes specific cytokine readouts [mRNA (*n* = 7 cytokines) or secretion (*n* = 12 cytokines) for 2D (*n* = 19 cytokines including both readouts) or 3D (*n* = 19 cytokines including both readouts) model systems] ranked based on mean reduction compared to LPS stimulation (black vertical line). Data were normalised to LPS or LPS with isotype, log-transformed (log_2_) and presented as mean ± SD. Mean responses that show at least a 20% inflammatory reduction (red vertical line) are framed, and the mean reduction (%) of the top ranked response is indicated. **(B, E)** Break down of top cytokine responses (shown as %) according to cohort, model system and readout. Highlighted in yellow are responses that show the highest proportion in each cohort. **(C, F)** Venn diagrams comparing common and unique top responsive cytokines to minocycline and IC14 among cohorts.

In terms of responses to minocycline, the HC and MCI cohorts displayed the highest number of top-ranked anti-inflammatory responses (14/38 and 15/38, respectively) compared to AD (5/38), with the top responses showing reductions of up to 64.5% (HC), 58.6% (MCI) and 49.7% (AD) of LPS-induced inflammation (**Fig. 4A; Supplementary data 1**). By breaking down the proportion of cytokines (*n* = 12 for secretion; *n* = 7 for mRNA) that showed efficacy in response to minocycline according to model system and cohort, we observed that in HC an even proportion of cytokines were effectively reduced at the secretion level, while more cytokines responded in 3D (85.7%) compared to 2D (57.1%) at the mRNA expression level (**Fig. 4B**). Similarly, MCI showed higher drug responsiveness in 2D for secreted cytokines (33.3%), and 3D for mRNA expression of cytokines (100%) (**Fig. 4B**). By contrast, in AD a higher proportion of cytokines showed reduction after drug treatment in 2D (16.7% for secretion, 28.6% for mRNA) compared to 3D (0% for secretion, 14.3% for mRNA) (**Fig. 4B**). This reveals differential minocycline efficacy depending on model system and cohort.

When comparing top-ranked cytokines among cohorts, common minocycline-responsive cytokines were identified in HC, MCI, and AD, including IL-6, IL-10, IL-1β and IL-18 (**Fig. 4C**). HC and MCI shared IL-8, TNF-α, TGF-β and IFN-γ, while IL-12p70 and IL-23 were found to be exclusively responsive in the MCI cohort (**Fig. 4C**).

In line with minocycline treatment, IC14 produced the highest number of anti-inflammatory responses in HC (15/38), followed by MCI (12/38) and AD (7/38) (**Fig. 4D; Supplementary data 2**). The highest-ranked responses in all cohorts displayed similar reduction levels (73.3% in HC, 72.4% in MCI and 70.8% in AD) (**Fig. 4D; Supplementary data 2**), suggesting that IC14 may have a more limited spectrum of action in AD but is equally efficacious across all cohorts. In both HC and AD, the 3D model showed a higher proportion of secreted cytokines (33.3% in HC and 16.7% in AD) that were reduced after IC14 treatment (**Fig. 4E**). In the MCI cohort no differences in the number of responsive cytokines were observed between 2D and 3D at the secretion level (**Fig. 4E**). At the mRNA expression level, the 2D model showed the highest proportion of responsive cytokines in all cohorts (100% in HC, 28.6% in MCI and 57.1% in AD) (**Fig. 4E**). IC14 targeted common cytokines across all cohorts, including IL-6, IL-1β, and IL-12p70. Shared targets between HC and MCI were IL-18, TNF-α and IFN-γ, while AD shared IL-8 with HC, and IL-10 and IL-23 with MCI. Additionally, unique responsive cytokines included IL-33 and TGF-β in HC, and MCP-1 in AD (**Fig. 4F**). Correlation analyses of minocycline and IC14 cytokine responses between model systems and readouts were not significant, likely due to low sample sizes (**Fig. S3, 4**).

This ranking analysis demonstrates that the performance of model systems and cytokine measurement readouts varies across cohorts, with notable differences observed in donors at distinct disease severity stages, such as MCI and AD. Furthermore, this approach facilitates the identification of responsive cytokines in a drug- and cohort-specific manner, potentially serving as biomarkers to guide future drug efficacy investigations.

### A novel approach to stratify treatment responders by accounting for disease state

Personalised drug testing approaches hold promise for improving the selection and stratification of treatment responders. Our analysis of 12 donors from all cohorts demonstrated that less than half exhibited anti-inflammatory responses to minocycline (5/12 in 2D and 4/12 in 3D), characterised by a minimum 20% reduction of LPS-induced inflammation (**Fig. 5A; Supplementary data 3**). The highest minocycline-mediated inflammatory reduction observed in the top donors was 39% and 50.9% in 2D and 3D, respectively. While the proportion of responders in the 2D model was greater in AD (67%) than HC and MCI (both 33%), in 3D minocycline was anti-inflammatory in HC and MCI (23% and 67%, respectively) but not in AD (0%) (**Fig. 5A**). Indeed, AD responders to minocycline (AD_1, AD_2) were identified only in the 2D model. In contrast, MCI responders were either common to both models (MCI_2) or unique to the 3D model (MCI_1) (**Fig. 5B**). Further donor response stratification by cytokine readout revealed that the main contribution in AD responders (AD_1, AD_2) was from mRNA expression in 2D, whereas in MCI (MCI_1, MCI_2) it came from both mRNA and secretion readouts (**Fig. S5A**).

**Fig. 5.**
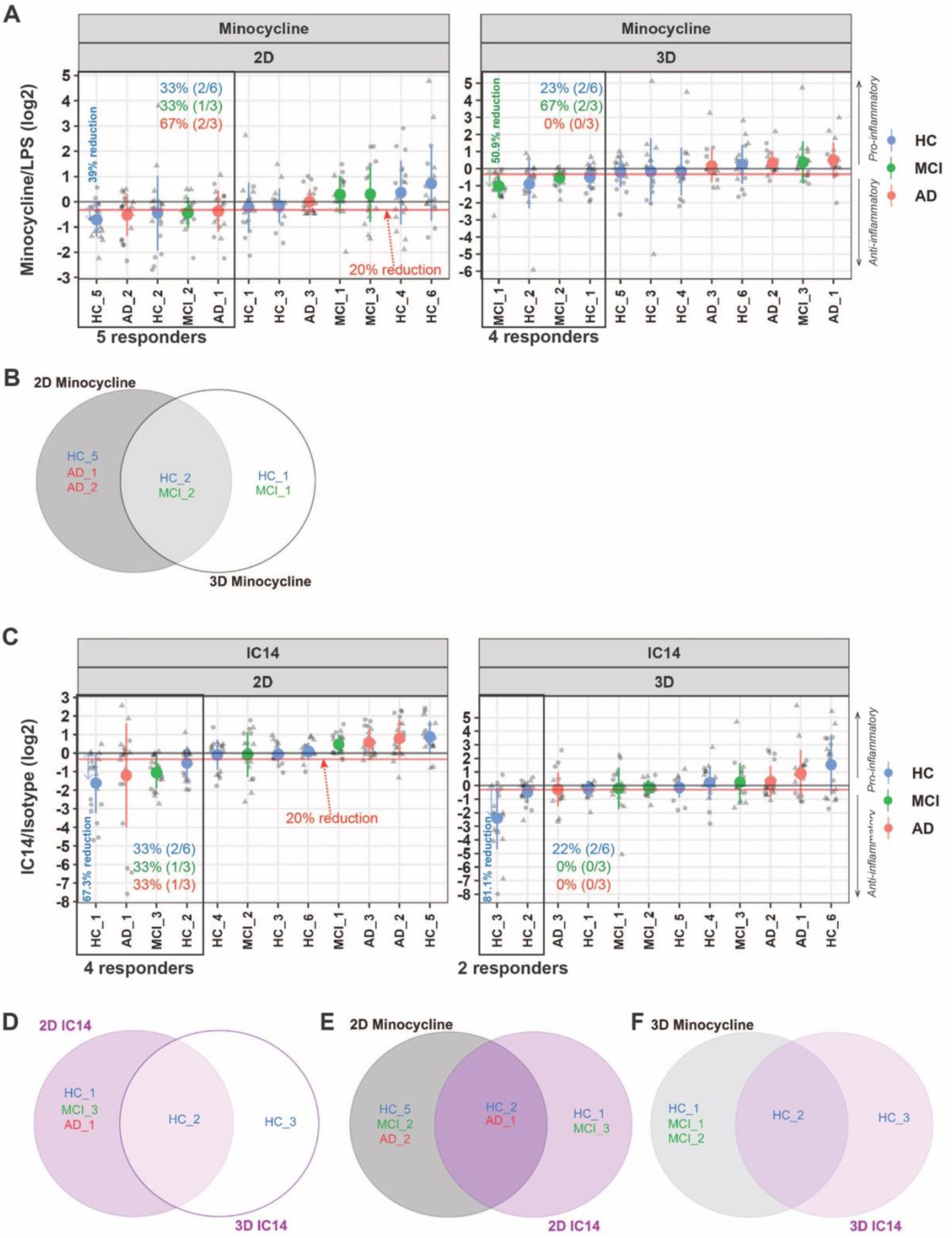
Ranking of donors to identify drug responders. **(A, C)** HC, MCI and AD donors ranked based on minocycline- or IC14-mediated reduction of their inflammatory profiles compared to LPS stimulation (horizontal black line). Data were normalised to LPS or LPS with isotype, log-transformed (log_2_) and presented as mean ± SD. Donors showing at least a 20% reduction of inflammation (horizontal red line) are framed, and the reduction (%) of the top ranked donor is indicated. **(B, D)** Venn diagrams comparing common and unique minocycline and IC14 responders between model systems. **(E, F)** Venn diagrams comparing common and unique responders between drugs in 2D and 3D.

Unlike minocycline, the number of responders to IC14 was different between models, with 4/12 in 2D and 2/12 in 3D, and a maximal inflammatory reduction of 67.3% in 2D and 81.1% in 3D (**Fig. 5C; Supplementary data 4**), indicating clear disparities in drug performance between models. In 2D, the proportion of responders (33%) was the same in all cohorts, while in 3D all responders were from the HC cohort (22%), with none coming from the MCI and AD cohorts (0% for both) (**Fig. 5C**). Interestingly, no MCI or AD donors were identified as common responders across models, suggesting that IC14’s effects in patient cohorts are specific to the model system used (**Fig. 5D**). For the patient responders (AD_1 and MCI_3) identified in the 2D model, the highest response to IC14 derived from the mRNA expression readout in AD, and more prominently from the secretion readout in MCI (**Fig. S5B**).

Differences between cohorts were observed regarding drug efficacy outcomes in each model system. HC donors were consistently responsive across model systems and drugs (**Fig. 5E-F**). However, for the patient cohorts responsiveness to both drugs was only found in the 2D model for AD (1 patient) (**Fig. 5E-F**), implying that the efficacy of anti-inflammatory drugs in MCI and AD patients is influenced by the type of model system used. Importantly, no AD responders to both minocycline and IC14 were discovered in the 3D model system (**Fig. 5F**). This highlights crucial differences in the suitability of these two drugs for AD.

Lastly, our investigation into the relationship between drug responses and disease risk factors, including age, *APOE* genotype, and brain amyloid-β burden, did not reveal any significant correlation (**Fig. S6, 7, 8**). Further research with larger sample sizes is warranted to validate these observations.

## Discussion

The lack of success in developing effective therapeutics for AD highlights the need for more robust drug development approaches. The genetic and biological heterogeneity among patients results in diverse pathophysiological profiles that can be effectively targeted through precision medicine strategies. Therefore, implementing precision drug development approaches may enhance the success of new therapeutics for AD (23). To improve the personalisation of drug discovery in AD, we have developed an *in vitro* drug testing pipeline that leverages the increased translational value of patient-specific MDMi cell models and 3D *in vitro* technology.

One of the main findings of our study is the ability to stratify participants based on the inflammatory cytokine profiles of their respective MDMi. Despite the small sample sizes (*n* = 3-6 donors) used to validate this pipeline, our hierarchical clustering and principal component analyses revealed a separation between profiles from healthy individuals and patients. Interestingly, this separation was more evident in 3D models and showed heterogeneity among HC, MCI and AD cohorts. This preliminary observation holds major implications for the current AD research landscape. Specifically, given our limited understanding of the molecular mechanisms driving disease onset and progression, the establishment of biomarker tools with robust predictive power remains challenging (24). Studying the inflammatory profiles of MDMi from patients with various disease severities and high-risk, asymptomatic individuals has the potential to inform improved diagnosis strategies and guide the implementation of preventative or therapeutic interventions at early stages. Nevertheless, future research is warranted to validate the sensitivity of MDMi inflammatory responses in reflecting inter-individual differences. For instance, in this study, HC donors had low non-*APOE*-associated genetic risk and were negative for brain amyloid-β deposition. A comparison of inflammatory profiles between these HC donors and another HC cohort with increased genetic risk and positive for brain amyloid-β deposition could provide valuable insights. Such differences in MDMi responses may provide an opportunity to guide personalised prevention trials aimed at minimising or delaying disease onset in high-risk individuals.

The pipeline presented in this study has the potential not only to discriminate asymptomatic individuals with different risk levels, but also to predict disease progression in patients before clinical symptoms manifest. Our observations revealed that in some cases, the inflammatory MDMi profiles of certain donors clustered more closely with individuals from other cohorts than their own. Although this could be attributed to high inter-individual variability, further investigation with larger cohorts (at least 20 donors) is warranted to determine if inter-cohort similarities reflect the conversion of HC to patients or the progression from mild to more severe disease stages. Elucidating this aspect would be crucial in defining the predictive ability of the developed pipeline and could guide better preclinical and clinical strategies to monitor disease onset and progression.

Recent studies in the field of cancer have highlighted the advantages of incorporating 3D models for precision drug development strategies (25–27). Similarly, in the field of neurodegenerative diseases there is a growing interest in using physiologically relevant 3D *in vitro* models as an alternative to animal models, which have low translational value (28–32). These 3D cell models have been used to demonstrate the successful pharmacological targeting of pathogenic alterations related to neurodegenerative diseases, including β- and γ-secretases in AD (33), the LRRK2 kinase in Parkinson’s disease (34), and the Src/c-Abl and mTOR pathways in amyotrophic lateral sclerosis (35). This demonstrates the potential of 3D models to enhance our mechanistic understanding of neurodegenerative diseases and validate candidate therapeutic compounds.

In our study, we have leveraged the personalised and clinically relevant features of MDMi to evaluate drug efficacy in 2D and 3D models. Our results revealed that both minocycline and IC14 reduced inflammation in MDMi from AD donors only when tested in 2D. In contrast, no patient donor showed drug responsiveness in 3D. This suggests a model-dependent mode of drug action in AD patient-derived cells and corroborates previous reports demonstrating the lack of clinical efficacy of minocycline in AD patients (16). To further validate the clinical translatability of the outcomes of our pipeline, testing other failed compounds in MDMi from larger patient cohorts could be a potential approach.

In contrast to AD, we observed that MDMi from most MCI donors showed responsiveness to minocycline in both 2D and 3D models, and IC14 in the 2D model alone. Similarly, MDMi from the HC cohort showed high drug response rates in both models. This supports the premise that treatment at early disease stages could provide a greater benefit than at later stages, where disease-related cellular and molecular alterations are irreversible (36). Likewise, modulating inflammation in asymptomatic individuals at high risk of developing dementia could even be more beneficial than in those with early disease symptoms. This reinforces the need for a tool that enables the early detection of individuals at risk based on their *in vitro* drug response profiles.

Despite the inter-individual heterogeneity, we were able to identify drug responders in each cohort, showing up to 81.1% of reduction in inflammation. This provides preliminary evidence that specific donors could potentially benefit from a treatment, providing a basis for recruiting more appropriately targeted subjects in clinical trials. Moreover, our findings suggest that a ‘one-size-fits-all’ treatment approach is unlikely to be effective for AD patients. Therefore, precision drug development strategies, including our *in vitro* drug testing pipeline, hold promise to increase the yield of effective disease-modifying therapies.

We performed a comprehensive evaluation of a range of inflammatory cytokines to assess drug efficacy in MDMi. Interestingly, we observed that cytokine modulation was drug- and cohort-specific, with higher numbers of drug responsive cytokines found in HC and MCI than the AD cohort. This indicates a potential difference in the inflammatory response and drug sensitivity among different disease stages. Specifically, commonly used cytokines to measure efficacy of anti-inflammatory drugs include IL-6 and IL-1β, which were modulated by minocycline and IC14 in all cohorts. However, our analysis revealed unique cytokines, namely MCP-1, IL-12p70, IFN-γ and IL-23, which were targeted by both drugs in the patient cohorts. Interestingly, MCP-1 and the IL-12/IFN-γ axis have been correlated with disease progression in clinical studies (37, 38), suggesting that these cytokine targets could be useful for directing future efficacy studies of new compounds in AD, such as IC14. Moreover, this provides a basis for designing new compounds that target specific inflammatory pathways that show high dysregulation in the context of disease.

In conclusion, we present an innovative *in vitro* drug testing pipeline that uses patient-derived MDMi cell models to assess the efficacy of microglia-targeted anti-inflammatory drugs with repurposing potential for AD. The implementation of this pipeline shows potential for enhancing patient stratification and responder selection strategies, thereby facilitating a more streamlined assessment of therapeutic drugs at the clinical level.

## Materials and Methods

### Study cohorts

Young healthy donors with no clinical symptoms (**Table 1**) were recruited at QIMR Berghofer Medical Research Institute (QIMRB-MRI), Queensland, Australia. Healthy control (HC), Mild cognitive impairment (MCI) and Alzheimer’s disease (AD) donors (**Table 2**) were recruited through the *Prospective Imaging Studying of Aging: Genes, Brain and Behaviour study* (PISA) at QIMRB-MRI. The recruitment strategy consisted of online cognitive assessments, genome-wide genotyping, multi-dimensional magnetic resonance imaging (MRI), amyloid positron emission tomography (PET) and neuropsychological testing (21). HC participants showed no cognitive alteration and were classified based on their genetic risk to develop AD. Those with low risk did not carry *APOE* ε4 and were in the lowest quintile of the AD PRS (polygenic risk score) no-*APOE* (*APOE* region excluded), which accounts for additional AD genetic risk variants. Experiments adhered to the ethical guidelines on human research outlined by the National Health and Medical Research Council of Australia (NHMRC) and have been approved by the Human Research Ethics Committees (HREC) of QIMRB-MRI and the University of Queensland. The donors’ informed consent was obtained before participation in the study.

### Isolation of peripheral blood mononuclear cells (PBMCs) from blood and generation of MDMi in 2D and 3D cultures

PBMCs were isolated from peripheral venous blood samples within 2h of blood withdrawal and frozen according to (6).

MDMi generation in 2D was performed as described in (6). Briefly, PBMCs were seeded onto Matrigel-coated (Corning, NY, USA) plates and monocytes (adherent cells) were cultured in serum-free RPMI-1640 GlutaMAX medium (Life Technologies, Grand Island, NY, USA) supplemented with 0.1 μg/mL of interleukin (IL)-34 (IL-34) (Lonza, Basel-Stadt, Switzerland), 0.01 μg/mL of granulocyte-macrophage colony-stimulating factor (GM-CSF) (Lonza, Basel-Stadt, Switzerland) and 1% (v/v) penicillin/streptomycin (Life Technologies, Grand Island, NY, USA) for 14 days.

MDMi generation in 3D was performed as described in (12). Briefly, PBMCs-derived monocytes were mixed with Matrigel, which had been previously diluted in culture medium at a 1:3 ratio. Matrigel-cell mixtures were then cultured in serum-free RPMI-1640 GlutaMAX medium containing 0.1 μg/mL IL-34 and 0.01 μg/mL GM-CSF for 30 days. Bright field images were captured using the Olympus CKX41 inverted microscope with 5.1MP CMOS digital camera.

### Treatment of MDMi with LPS and drugs

LPS (O55:B5, Sigma-Aldrich, MO, USA) was dissolved in RPMI-1640 GlutaMAX medium and stored at -20°C, as per manufacturer’s instructions. For MDMi treatment, LPS was added to 2D and 3D MDMi at a concentration of 10 ng/mL for 24h. Treatments with the FDA-approved drugs minocycline and IC14 were performed in 2D MDMi after 14 days of culture, and in 3D MDMi after 30 days of culture. Drug concentrations were optimised in 2D cultures, as 3D cultures withstand higher drug concentrations. Minocycline (#252859, Sigma-Aldrich, MO, USA) was reconstituted in DMSO and used at 1 μM for 24h. IC14 (kindly provided by Implicit Bioscience) was used in at 0.05 ng/mL for 24h. DMSO was used as vehicle control for minocycline treatments. An IgG4 antibody (#403702, Australian Biosearch, WA, Australia) was used as isotype control for IC14 treatment. To assess the capacity of minocycline and IC14 to reduce LPS-induced inflammation in 2D and 3D MDMi, drugs or the respective vehicle controls were added for 1h in 2D cultures, or 2h in 3D cultures, prior to LPS exposure. Following drug pre-treatment, cultures were stimulated with 10 ng/mL LPS. Cultures were harvested for downstream analysis after 24h of LPS incubation.

### Statistical analysis

For cohort-based analyses, GraphPad Prism software version 9 (GraphPad Software, CA, USA) was used. Data were presented as bar graphs with mean ± SEM or SD. Cytokine responses from young donors to increasing drug concentrations were averaged and compared using repeated-measures one-way ANOVA. To measure drug efficacy in the cohorts, drug responses were calculated relative to LPS-stimulated responses and log transformed and efficacy was tested using one-sample *t*-test against a null hypothesis (log(relative change) = 0).

For donor-specific analyses, R package (version 4.1.0) was used. Cytokine responses were log-transformed. A Kruskal-Wallis test was used to examine differences between cohorts in drug responses. Spearman correlation was used for correlation analyses. For multivariate analyses, missing values were imputed with the mean of the log-transformed relevant cytokine values and data were centered and scaled. Principal component analysis (PCA) was visualised using plots where enclosing ellipses were estimated using Khachivan algorithm. Hierarchical clustering was preformed using Euclidian distance and complete linkage clustering, and visualised using heatmaps. For ranking analyses, mean cytokine and donor responses were ranked based on geometric mean fold change. Drug-mediated reduction of inflammation was presented as a percentage change from LPS-treated responses.

## Additional Materials and Methods

Additional methods describing immunocytochemistry, confocal imaging and cytokine RNA and protein secretion analyses are available in the SI Appendix.

## Author Contributions

C.C-L., R.S., A.R.W. and H.Q. conceived and designed the study. C.C-L. and H.Q. performed experiments and interpreted data. S.O. performed donor-specific statistical analyses and correlations. G.L.R. and M.W.A. kindly provided IC14. C.C-L. wrote the manuscript and made figures. R.S., A.R.W. and H.Q. interpreted data and provided critical feedback in reviewing and editing. All authors reviewed and commented on the final version of the manuscript.

## Acknowledgments

We thank Dr Gunter Hartel for his invaluable advice on data analysis. We acknowledge Sun Yifan Emily and Natalie Garden for their assistance with donor blood collection, and the coordinators of the PISA study Professor Michael Breakspear, A/Professor Michelle Lupton, Dr Christine Guo, Jessica Adsett and Madeline Wood. We appreciate the generosity of the QIMRB-MRI volunteers and PISA study participants, who have kindly donated blood to this project. We also thank the QIMRB-MRI Scientific Services and Sample Processing teams for their support with experimental procedures and collection of samples.

